# Challenges in quantifying functional redundancy and selection in microbial communities

**DOI:** 10.1101/2024.03.26.586891

**Authors:** Po-Yi Ho, Kerwyn Casey Huang

## Abstract

Microbiomes can exhibit large variations in species abundances but high reproducibility in abundances of functional units, an observation often considered evidence for functional redundancy. Based on such reduction in functional variability, selection is hypothesized to act on functional units in these ecosystems. However, the link between functional redundancy and selection remains unclear. Here, we show that reduction in functional variability does not always imply selection on functional profiles. We propose empirical null models to account for the confounding effects of statistical averaging and bias toward environment-independent beneficial functions. We apply our models to existing data sets, and find that the abundances of metabolic groups within microbial communities from bromeliad foliage do not exhibit any evidence of the previously hypothesized functional selection. By contrast, communities of soil bacteria or human gut commensals grown *in vitro* are selected for metabolic capabilities. By separating the effects of averaging and functional bias on functional variability, we find that the appearance of functional selection in gut microbiome samples from the Human Microbiome Project is artifactual, and that there is no evidence of selection for any molecular function represented by KEGG orthology. These concepts articulate a basic framework for quantifying functional redundancy and selection, advancing our understanding of the mapping between microbiome taxonomy and function.

## INTRODUCTION

Functional redundancy is the concept that multiple species within a microbial community can share the same function, and is often associated with the general idea that function rather than taxonomy determines the assembly of microbial communities^1-4^ via selection for functional units^5^. Functional redundancy has been hypothesized to underlie the robust functioning of wide-ranging phenomena, including ocean microbiome-associated biogeochemical cycles^6^, human gut microbiome variability^7,8^, and wastewater treatment by activated sludge^9^, and is often inferred empirically through the observation that a high degree of variation in species abundances appears relatively constant when converted to abundances of functional units (i.e., groups of species that share metabolic capabilities, gene families, or other traits^10-16^). However, this reduction in variability does not imply functional redundancy or selection for functional units if other factors can also reduce variability. Here, we clarify the link between functional variability and selection for functional units. We identify potential confounders that generate apparent selection, develop models to disentangle their effects, and apply our models to find potential units of functional selection across diverse microbiomes. Our results provide a new lens to interpret microbiome function.

## RESULTS

### Reduced functional variability does not imply functional selection

An intuitive hypothesis is that if selection is acting on functional units, then the variability in functional abundances should be less than that of taxonomic abundances. However, the converse is not true because functional variability can be lower than taxonomic variability due to factors other than selection on functional abundances. Consider a collection of communities {𝒞} with abundance 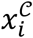 and 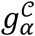 for taxon *i* and function *α*, respectively. Taxonomic compositions 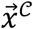 and functional profiles 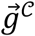 are related by the matrix *G* that describes the gene content of each species such that

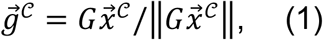

Where *G*_*αi*_ *=* 1 denotes that taxon *i* contains function *α* and *G*_*αi*_ *=* 0 otherwise, and ‖ ‖ denotes the sum over all elements. Here, for simplicity we did not consider gene copy number, which can be straightforwardly incorporated into the analyses described below. Eq. 1 assumes that the taxonomy-function map is linear and conserved across communities, which have recently been found to approximately hold across various ecosystems^17^. Transformation according to Eq. 1 can also describe the process of grouping species by other signatures such as phylogeny, motivated by the assumption that species from the same coarse taxon are more likely to share common functions. In any case, for diverse communities, this transformation involves summing over many taxa, and hence by the central limit theorem, the abundance of a function will tend to scale with the number of taxa that harbors it regardless of variability in taxonomic abundances. Consequently, the functional distance 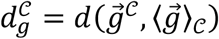 from a given composition 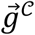 to the mean composition across communities 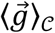 will typically be less than the corresponding taxonomic distance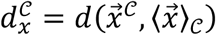, where *d* is any dissimilarity metric such as Bray-Curtis. At the ecosystem level, the average functional distance from the mean profile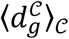, a measure of functional variability for a collection of communities, will also typically be less than the average taxonomic distance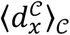 .

To demonstrate this statistical averaging effect, we randomly generated a gene content matrix and a set of 200 communities *in silico* with 10 functions and 30 species. Relative abundances were drawn from the abundance distributions of Human Microbiome Project (HMP) stool samples to capture a natural degree of taxonomic variability (**Methods, Fig. 1a**). Functional compositions were obtained via matrix multiplication as in Eq. 1. Species abundances were sampled in an uncorrelated manner, and no selection was applied to functional compositions. Nonetheless, the statistical averaging effects described above alone led to substantial functional similarity (**Fig. 1b**), reminiscent of previous claims of functional selection^7,12^. Thus, smaller functional variability does not by itself imply selection on functional units.

**Figure 1:**
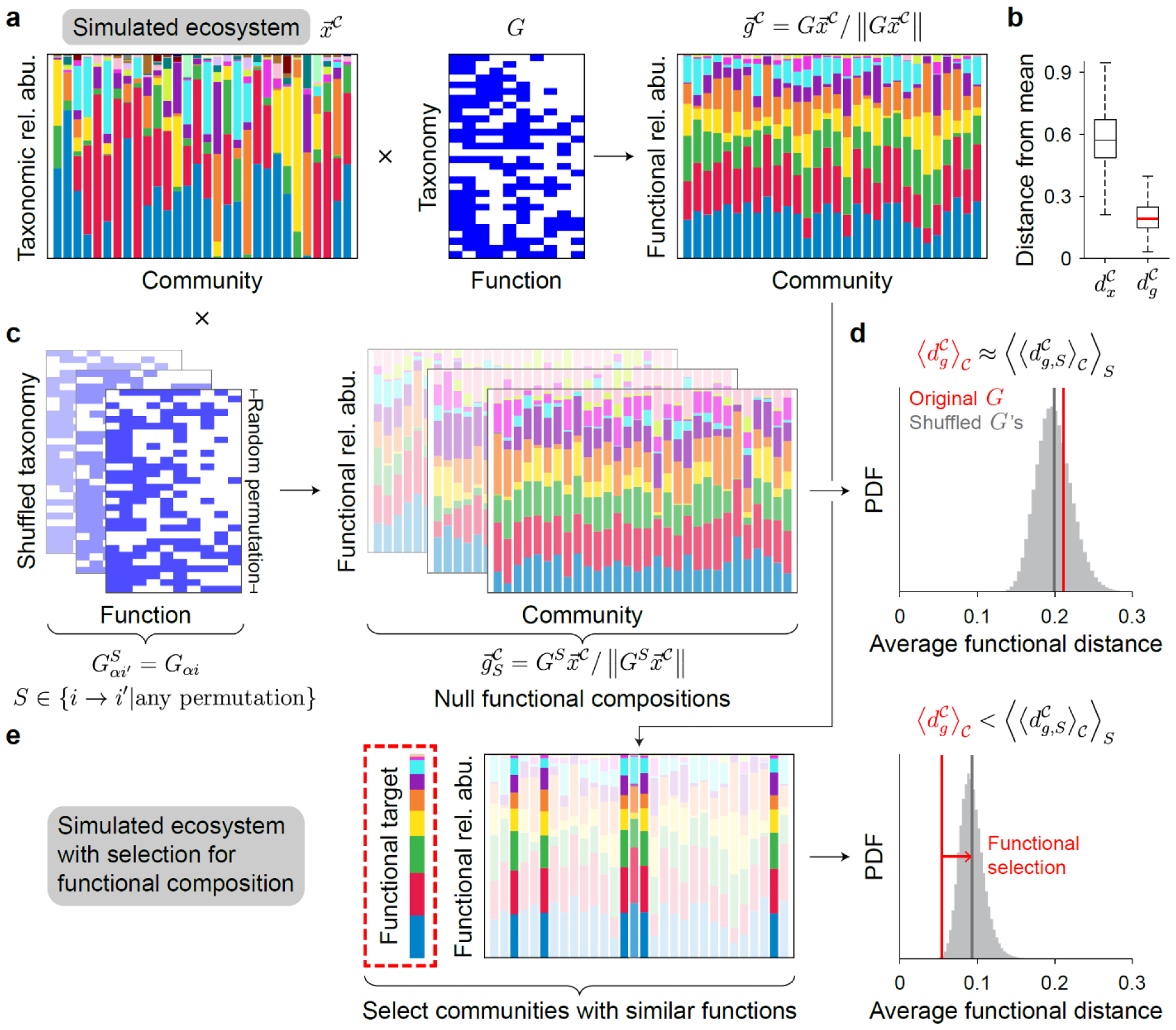
Functional similarity does not imply functional selection since statistical averaging effects also increase functional similarity. a) Functional compositions are obtained from taxonomic compositions via matrix multiplication, illustrated using a simulated ecosystem with 30 species, 10 functions, and 200 communities (a random subset is shown). Species relative abundances 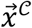 in communities 𝒞 were randomly drawn from the species abundance distributions of HMP stool samples (**Methods**). The gene content matrix *G* was generated randomly (**Methods**). The functional relative abundance profiles 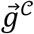 were calculated via Eq. 1. b) Without any selection on functional units, statistical averaging effects alone led to substantial functional similarity. Shown are the distributions of Bray-Curtis dissimilarity metric for each community relative to the composition of the mean community for species 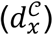 and functional 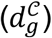 abundances. The central mark, box, whiskers, and plus signs denote the median, the 25^th^ and 75^th^ percentile, and the extremes, respectively. c) Null functional compositions obtained from shuffled gene content matrices *G*^*S*^ were also more similar than taxonomic compositions. *S* indicates a random permutation of the taxa labels {*i*}. Shown are three examples of random shuffling. d) The random communities were not more functionally similar compared to the null functional compositions. The distribution of the distance from the mean community averaged across communities for 2×10^5^ random shuffles (gray) is compared to the average distance from the mean for the original gene content matrix (red). e) Functional selection was imposed on the ensemble of randomly generated communities in (a) by selecting the top 1% of communities whose functional abundances were most similar to a target functional composition. In this example, the mean functional composition of the original ensemble was set as the target. The average functional distance for the original gene content matrix is now significantly smaller than the average distance after shuffling.

### An empirical null model to quantify functional selection

Instead, the extent of functional selection must be determined by comparing against null models that account for statistical averaging. A natural option is to substitute for *G* in Eq. 1 with another matrix *G*^*S*^ obtained by shuffling the taxa {*i*} by a random permutation *S*^8,18^ (**Fig. 1c**):

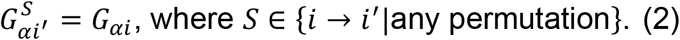

Such random shuffling preserves the effect of statistical averaging under the same functional prevalences as the original gene content matrix, but scrambles correlations between taxa and functions. If the decrease in functional variability (**Fig. 1b**) is due solely to statistical averaging, then the functional variability obtained after shuffling 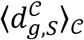 will be approximately the same as the original variability when averaged over many random shuffles 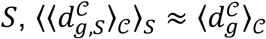 (**Fig. 1d**).

By contrast, if the ecosystem is selecting for a specific functional profile, correlations between taxa and functions should contribute to decrease functional variability. In this case, shuffling eliminates those correlations and the functional variability after shuffling should be greater than before,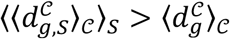 . To demonstrate this effect, we selected, from among the ensemble of randomly generated communities described above, the top 1% of communities whose functional compositions were most similar to a target profile (**Methods, Fig. 1e**), thus simulating the effect of selection for a particular functional profile. As expected, this selection decreased functional variability in a manner that was dependent on the correlations between taxa and function encoded by the original gene content matrix. Indeed, comparison with randomly shuffled matrices showed that 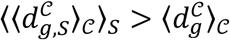, and the difference between these two quantities represents the extent of functional selection (**Fig. 1e**). This empirical approach for constructing null models can be applied to analyze various ecosystems without making additional assumptions about the distributions of taxonomic and functional abundances and the correlations between them.

### Identifying the extent of functional selection in diverse microbiomes

We applied this null model to existing data sets to determine whether reduced functional variability is due to functional selection. We focused on metabolic capability because if the community dynamics in an ecosystem are driven by resource competition, then species abundances will be determined largely by metabolic capabilities rather than other species-specific properties. First, we investigated microbial communities inside the foliage tanks of bromeliads, which (as previously shown^12^) exhibited decreased variability when taxonomy was converted via Eq. 1 to metabolic gene groups including methanogenesis, fermentation, and chemoheterotrophy (**Methods, Fig. 2a**). Comparison against random shuffles revealed that the observed functional variability was not significantly smaller than that after shuffling; in fact, functional variability increased (**Fig. 2b**), suggesting that the reduction in the variability of custom metabolic gene groups arose from statistical averaging effects and not functional selection for these metabolic gene groups.

**Figure 2:**
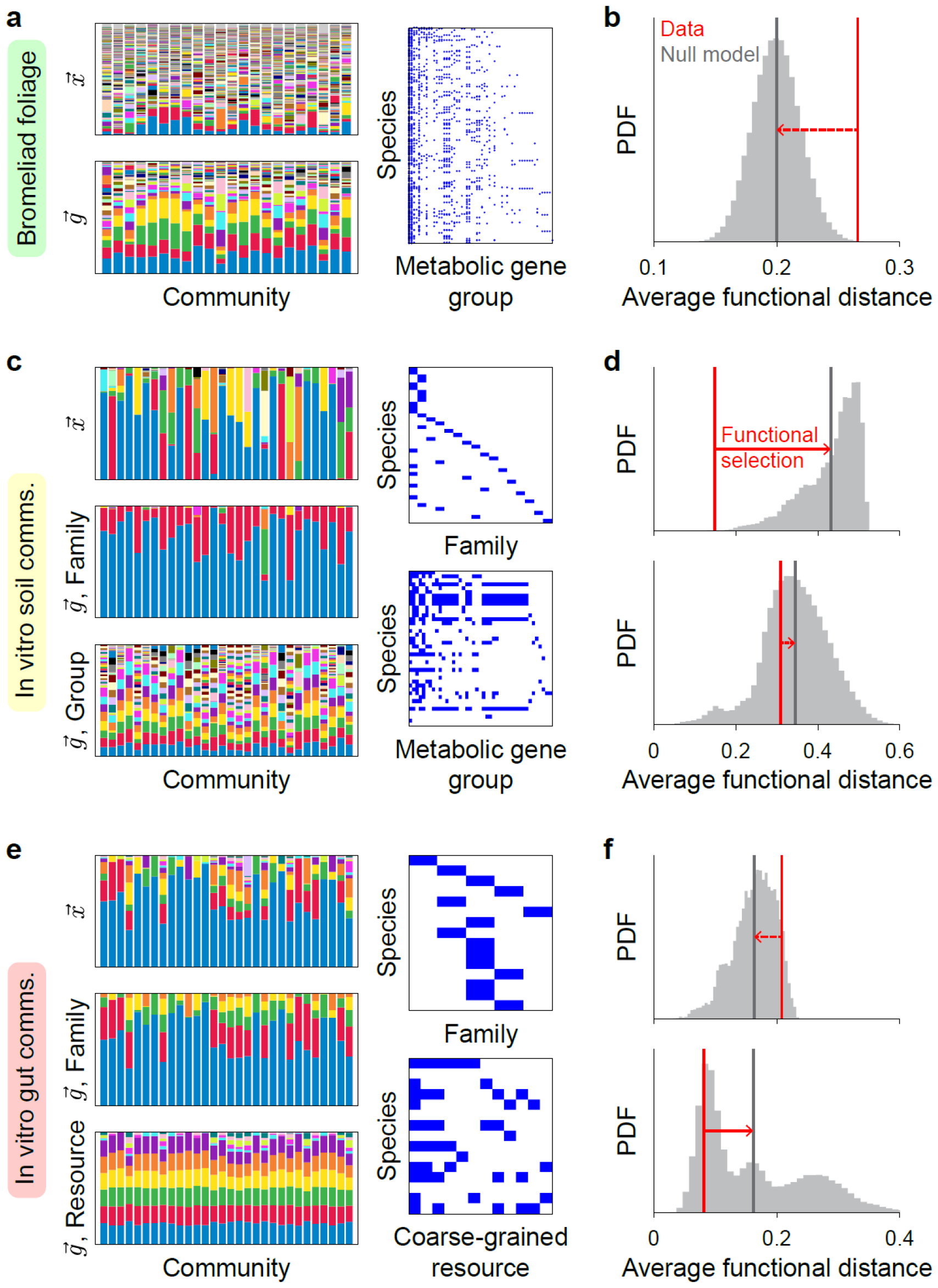
Comparison with random shuffles reveals the extent of functional selection in various microbiotas. a) Species abundances, functional abundances, and the taxonomy-function map of microbial communities found in bromeliad foliage tanks^12^. All data were obtained from previous studies (**Methods**). Metabolic gene groups include aerobic chemoheterotrophy, fermentation, methanogenesis, and other common classifications. b) For the bromeliad communities in (a), the average functional distance from the mean community (red) was not lower than those obtained from random shuffling (gray). c) Same as (a) but for soil- and leaf-derived communities grown *in vitro* in minimal medium supplemented with glucose as the single carbon source^14^ (**Methods**). Prominent families include Enterobacteriaceae (blue) and Pseudomonadaceae (red). d) Same as (b) but for the communities in (c). Here, the average functional distance was significantly lower than those obtained from random shuffling. e) Same as (a,c) but for gut bacterial communities assembled *in vitro* in complex medium^19^ (**Methods**). The communities were assembled from 15 species spanning 6 families. The coarse-grained resource utilization profiles were determined from metabolomics data for each species grown individually^19^. f) Same as (b,d) but for the communities in (e) using taxonomic families (above) or coarse-grained resources obtained in a previous study^19^ as the functional units.

To search for an example of functional selection, we investigated a simpler ecosystem of soil- and leaf-derived *in vitro* communities grown in minimal media^14^. These communities consisted mainly of species from two taxonomic families, Enterobacteriaceae and Pseudomonadaceae; in minimal media with glucose as the sole carbon source, many of the Enterobacteriaceae species consumed glucose and secreted acetate, which were then utilized by the Pseudomonadaceae species^16^. Thus, taxonomy is tightly linked to metabolic capability in this ecosystem, and consequently, the communities exhibited significantly decreased variability when species were grouped by taxonomic family^14^. In contrast to the bromeliad communities, variability was small even when compared against random shuffles of the species-family map (**Methods, Fig. 2c,d**). This result is not generalizable to all taxonomy-function maps, since converting species abundances to the metabolic gene groups used in the bromeliad example did not lower functional variability compared to random shuffles (**Fig. 2c,d**). These findings suggest that selection for specific metabolic capabilities indeed underlies the assembly of these communities.

Next, we tested selection for metabolic capability in assemblies of human gut bacteria *in vitro*^19^, a relatively more complex ecosystem in terms of taxonomy and nutrients. In these communities, grouping species into families did not lead to reduced variability relative to random shuffling (**Methods, Fig. 2e,f**). This failure of family-level grouping to reduce variability is consistent with the observation that phylogenetic relatedness does not predict growth behaviors in this set of species^20^. Instead, in a previous study, we used coarse-grained metabolomic profiles to identify species with similar metabolic capacity^19^, and converting species abundances to metabolic capability via these groupings identified functional selection (**Methods, Fig. 2e,f**). These results are broadly consistent with multiple studies showing that metabolism is not always correlated with phylogeny^21^ but is a major driver of *in vitro* assembly^19,22,23^. More broadly, our simulations and analyses highlight that claims of selection for functional profiles and algorithms for finding such profiles must be tested against null models.

### Functional bias can reduce functional variability without ecosystem-specific selection

Next, we analyzed HMP stool samples in terms of molecular functions represented by KEGG orthologs (KOs). These communities exhibited reduced variability in the abundances of core pathways such as glycolysis and ATP synthesis^7^, even when compared with random shuffles of the taxonomy-function map (**Methods, Fig. 3a**). However, many of these core pathways likely confer fitness benefits regardless of any selection collectively imposed by the ecosystem, and we hypothesized that these benefits could reduce functional variability independent of ecosystem-specific selection for communities that match a target profile of these core pathways. Indeed, taxa with these beneficial functions typically exhibited higher abundance than those without (**Fig. 3b**), a property that we term functional bias. Unlike selection for a target composition of multiple functions, bias is a property of an individual function and can arise independently of ecosystem-specific selection for functional composition, as we demonstrate below. Whereas selecting communities based on functional composition directly affects functional variability, as we showed above (**Fig. 1e**), we reasoned that functional bias can affect functional variability *indirectly* through species abundances.

**Figure 3:**
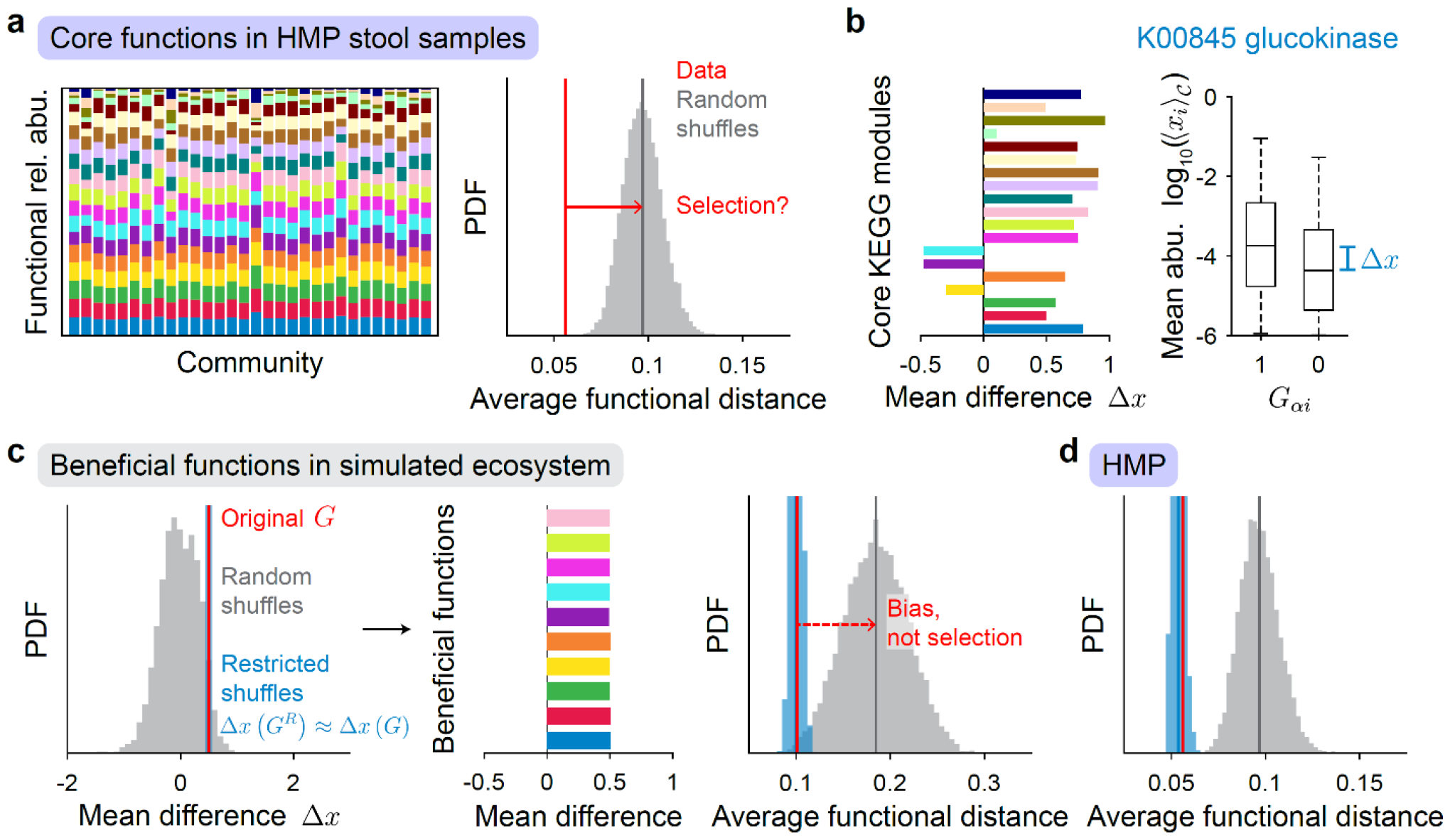
Beneficial functions tend to decrease functional variability without selection on functional profiles. a) HMP stool samples exhibited reduced variability in the abundances of core functions (**Methods**). We selected 19 KEGG orthologs that are the primary entry in each of 19 KEGG modules, representing central carbohydrate metabolism, amino acid metabolism, ATP synthesis, and other core functions. b) A beneficial function is a function for which the difference Δ*x* between the mean abundance of species with and without the function is positive. Of the 19 KOs analyzed here, 16 are beneficial. c) Restricted shuffles, unlike completely random shuffles, approximately preserve the mean difference Δ*x* (**Methods**). Functional bias was introduced independently of ecosystem-dependent selection by generating beneficial functions using restricted shuffles. In the example shown, 10 beneficial functions with target Δ*x =* 0.5 were generated. The resulting taxonomy-function map led to reduced functional variability compared to random shuffles, but not restricted shuffles. d) Restricted shuffles applied to analyze the functional profiles of the HMP stool samples in (a) demonstrates a similar reduction in functional variability compared to random but not restricted shuffles, as in (c).

To demonstrate the effects of functional bias, we developed an algorithm to generate a function *α* defined by species combinations *G*_*αi*_ that approximately exhibits a pre-specified mean difference

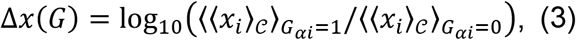

where ⟨ ⟩_G*αi=*1,0_ denotes averaging over all species *i* with or without the function, respectively (**Methods, Fig. 3c**). In other words, the algorithm enforces that function *α* confers a mean log abundance difference Δ*x*. Using this algorithm, we generated 10 beneficial functions with target mean difference Δ*x =* 0.5 and found that the gene content matrix obtained by combining these 10 functions led to reduced functional variability versus random shuffles (**Methods, Fig. 3c**).

Importantly, we did not select or remove from the ensemble any of the randomly generated communities as we did in **Fig. 1e** to demonstrate the effects of selection on functional profiles. This result demonstrates that functional bias can lead to apparent functional selection when evaluated using completely random shuffling, which abolishes all correlations between taxa and function regardless of whether they arise from ecosystem-dependent selection or bias. To separate the distinct contributions of these two factors, we performed restricted shuffles *R* for function *α* that preserve the total number of species with function *α* while also approximately maintaining the mean difference:

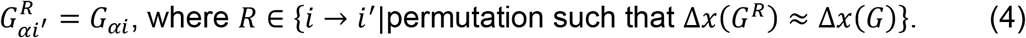

Restricted shuffles retain correlations between taxa and function that mimic the indirect effects of functional bias (**Fig. 3c**). Applying restricted shuffles to the HMP communities, we found that they did not exhibit reduced variability in the abundances of core pathways (**Fig. 3d**), suggesting that these communities were not selected for their profiles of the core pathways.

### No selection for functional orthologs in human gut microbiome

While the HMP communities as a whole did not exhibit functional selection for core pathways, it remained possible that specific functions could exhibit signatures of ecosystem-dependent selection. To comprehensively test this hypothesis, we considered each function individually. As a case study, we analyzed a highly prevalent KO, K00845 (glucokinase), which exhibited a relatively large mean difference of Δ*x*(*G*) = 0.79. We evaluated the abundance of each function *α* without normalizing by the total abundance, 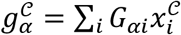 and computed the coefficient of variation (CV, the standard deviation divided by the mean) as a measure of variability. For K00845, the CV of its functional abundance across communities was significantly lower compared to the distribution of CVs obtained after completely random shuffling (**Fig. 4a**): the *z*-score was -3.2 (*p* = 6.1×10^-4^), suggesting (misleadingly) that this function is selected for low variability. However, the *z*-score relative to the distribution of CVs obtained by restricted shuffling was only -1.2 (*p* = 0.11). Thus, functional variability was significantly lower compared to unrestricted but not restricted shuffles, implying that functional bias rather than selection drives the overall reduction in the functional variability of this KO.

**Figure 4:**
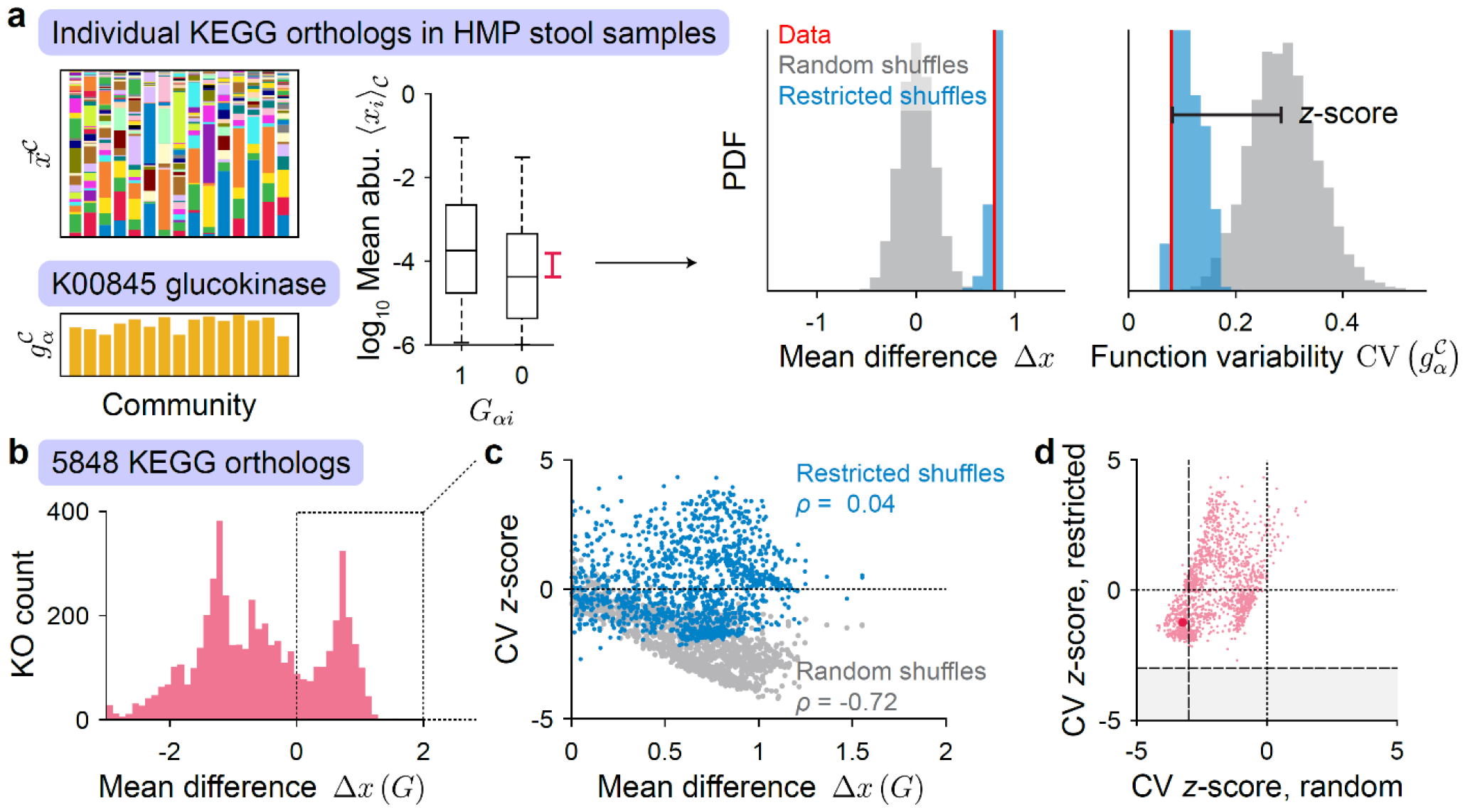
Separating functional bias via restricted shuffling reveals that the human gut microbiome is not selected for any KEGG ortholog. a) KEGG ortholog K00845 as an example of a beneficial function in HMP stool samples (**Methods**). Random shuffles (gray), but not restricted shuffles (blue), typically led to higher functional variability, as measured by the CV of the abundance 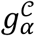 of the beneficial function *α* . The CV *z*-score (black bar) is the difference between the functional variability and the average functional variability over random or restricted shuffles, normalized by the standard deviation across shuffles. b) Distribution of Δ*x* for all analyzed KOs, showing that a substantial fraction exhibited positive Δ*x*. c) For KOs with positive Δ*x*, the CV *z*-score was negatively correlated with the mean difference when calculated relative to random shuffles (gray), but not relative to restricted shuffles (blue). d) Restricted shuffles reduced the magnitude of CV *z*-scores. Of the KOs with positive Δ*x*, none exhibited significantly reduced functional variability (CV *z*-score <-3) when compared to restricted shuffles (gray region). The highlighted point denotes the KO in (c)

Of the 5,848 KOs evaluated, 1,554 (27%) exhibited functional bias with Δ*x*(*G*) > 0 (**Fig. 4b**). Among these KOs, there was a significant negative correlation between the mean difference Δ*x*(*G*) and the CV *z*-score relative to unrestricted shuffles (Pearson’s correlation coefficient *r* = -0.72, *p* = 1.6×10^-250^, **Fig. 4c**), indicating that the effects of functional bias were prevalent and tended to reduce functional variability. By contrast, there was not a significant correlation when the *z*-score was calculated using restricted shuffles (*r* = 0.04, *p* = 0.15, **Fig. 4c**). In fact, none of the 354 KOs that had a significant z-score (<-3) relative to unrestricted shuffles had a significant *z*-score when compared to restricted shuffles (**Fig. 4d**). Thus, there was no evidence that stool samples from the HMP were selected for any KO. Whether the human gut microbiome is under functional selection, and if so, the functional unit under selection remain outstanding questions.

## DISCUSSION

Here, we showed that statistical averaging and functional bias are confounding factors when assessing functional redundancy and selection, and proposed empirical null models to disentangle their effects. There are likely additional factors that can further confound analysis of functional variability. Thus, it will be important to continue to develop the appropriate null models and incorporate them into unsupervised identification of selected units^24,25^. A new concept emerging from our analyses is the possibility for increased, rather than reduced, functional variability relative to null models (**Fig. 2a**), which will be interesting to understand as a contrast to functional redundancy. An important limitation is that gene abundance may not be representative of the realized capacity of a function, and null models must also be developed to assess functional redundancy in transcriptomics, proteomics, metabolomics, and other functional data. These models must accommodate potential nonlinearity in the taxonomy-function map that might not easily described by a gene content matrix as in Eq. 1. Although outstanding questions remain, further work based on the concepts proposed here will ultimately shed light on the unit of selection in microbial communities and provide new tools to control microbiome function.

## METHODS

### Data sets

All data sets analyzed were obtained from existing studies. All code and associated data will be available in a public repository on publication.

#### Bromeliads

Data were obtained from Ref. ^12^, in which detritus from the bottom of bromeliad foliage tanks was collected and sequenced via 16S rRNA gene amplicon sequencing. Operational taxonomic units (OTUs) were identified from sequencing data and associated with one or more metabolic functions based on the functional annotations of the prokaryotic taxa (FAPROTAX) data base^6^. In FAPROTAX, a taxon is assigned a function, such as aerobic chemoheterotrophy, fermentation, or methanogenesis, if all cultured species within that taxon are known to exhibit that function based on extensive literature searches.

#### Human Microbiome Project

Data were obtained from Ref. ^7^, involving 142 stool samples from the HMP phase 1 study, which consisted of 82 healthy subjects, some of which were sampled multiple times (https://hmpdacc.org/hmsmcp2/). Species relative abundances were obtained from metagenomic sequencing data as described in the study. The gene content network was obtained from Ref. ^8^, which associated 796 species with 7,015 KEGG orthologs (KOs) using reference genomes in the Integrated Microbial Genomes & Microbiomes-Human Microbiome Project (IMG/M-HMP) database. Each KO is a group of genes representing functional orthologs. For species that have multiple reference genomes in the database, a representative genome was randomly chosen. We analyzed the 131 samples in which the gene content network contained representative genomes for species whose total relative abundance was >80%; these samples contained 5,848 KOs.

#### Soil- and leaf-derived *in vitro* communities

Data were obtained from Ref. ^14^, in which communities were inoculated into M9 minimal media supplemented with a single carbon source and passaged via serial dilution until an ecological steady state was reached in which the community composition in subsequent passages remained approximately the same. We analyzed the 16S sequencing data at the end of serial passaging for 96 communities derived from 12 inocula with 8 replicates each, all grown in M9 with glucose.

#### Communities of gut commensals grown *in vitro*

Data were obtained from Ref. ^19^, in which 15 human gut bacterial species from 6 families were assembled and grown *in vitro* in the complex growth medium Brain Heart Infusion (BHI): Enterobacteriaceae (*Escherichia fergusonii*), Enterococcaceae (*Enterococcus hirae, Enterococcus faecium, Enterococcus faecalis*), Bacteroidaceae (*Bacteroides thetaiotaomicron, Bacteroides fragilis, Bacteroides uniformis*), Lachnospiraceae (*Clostridium symbiosum, Clostridium clostridioforme, Clostridium scindens, Clostridium hylemonae, Clostridium hathewayi, Blautia producta*), Oscillospiraceae (*Flavonifractor plautii*) and Porphyromonadaceae (*Parabacteroides distasonis*). The coarse-grained resource utilization structure was obtained from metabolomics data for each species grown individually as described in Ref. ^19^. Communities were passaged via serial dilution until they reached an ecological steady state. We analyzed the 16S sequencing data at the end of serial passaging for 80 communities of ≥3 species.

#### Simulated random communities

We generated random ecosystems to develop null models that account for statistical averaging and functional bias. To generate a random gene content matrix, each matrix element *G*_*iα*_ for species *i* and function *α* was set to 1 with probability *P*_*α*_, the prevalence of function *α*, and 0 otherwise. *P*_*α*_ was evenly spaced between 0.2 and 0.8 across *α*, mimicking the observed range of prevalences. To generate random communities, the abundance 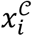 of species *i* in community 𝒞 was randomly drawn from the distribution of abundances for a random species in a real data set and then normalized for each community. This approach based on empirical data avoids the need for additional assumptions about species abundance distributions. To generate a random beneficial function *α*, the simulated annealing algorithm described below was applied for given values of prevalence *P*_*α*_ and mean difference Δ*x*(*G*_*α*_). The numbers of species, functions, and communities, as well as *P*_*α*_ and Δ*x*(*G*_*α*_) were set to typical values provided in the relevant figure legends.

#### Metropolis-Hastings simulated annealing algorithm for restricted shuffles

The generation of restricted shuffles that preserve the target mean difference Δ*x*(*G*) requires an efficient sampling algorithm over all possible permutations. To accomplish this goal, we implemented the Metropolis-Hasting simulated annealing algorithm from statistical physics. The algorithm began with a random permutation *R* over all taxa. At each iteration, the algorithm picked a taxon that contained the function being considered and another that did not. To determine whether to swap these two taxa to form permutation *R*′, the energy of permutation *R*′ was calculated as the absolute value of the difference between the mean difference (*Δx*) under the permutation and the target mean difference: *E*(*G*^*R*^′) *=* |*Δx*(*G*^*R*^′) − *Δx*(*G*)| . The swapped permutation *R*′ was kept with probability exp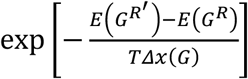 ; otherwise, the original was kept. Here, *T* denotes a temperature parameter that controls the acceptance probability. This process was repeated for 100 iterations at each temperature in the simulated annealing schedule *T =* {1, 0.1, 0.01, 0.005, 0.001, 0.0001}. The algorithm concluded at the end of the schedule, resulting in one restricted shuffle that approximately preserves the target mean difference.

## ACKNOWLEDGEMENTS

We thank Benjamin Good, Karna Gowda, and members of the Huang lab for helpful discussions. This work was funded by the Stanford School of Medicine Dean’s Postdoctoral Fellowship (to P.H.), NIH Postdoctoral Fellowship F32 GM143859 (to P.H.), NSF Awards EF-2125383 and IOS-2032985 (to K.C.H.), and NIH Awards R01 AI147023 and RM1 GM135102 (to K.C.H.). K.C.H. is a Chan Zuckerberg Biohub Investigator.

